# Nano immunoconjugates crossing blood-brain barrier activate local brain tumor immune system for glioma treatment

**DOI:** 10.1101/466508

**Authors:** Anna Galstyan, Antonella Chiechi, Alan J. Korman, Tao Sun, Liron L. Israel, Oliver Braubach, Rameshwar Patil, Ekaterina Shatalova, Vladimir A. Ljubimov, Janet Markman, Zachary Grodzinski, Keith L. Black, Manuel L. Penichet, Eggehard Holler, Alexander V. Ljubimov, Hui Ding, Julia Y. Ljubimova

## Abstract

Treatment of brain gliomas with checkpoint inhibitor antibodies to cytotoxic T-lymphocyte-associated antigen 4 (a-CTLA-4) and programmed cell death-1 (a-PD-1) was largely unsuccessful due to their inability to cross the blood-brain barrier (BBB). We describe a new generation of nano immunoconjugates (NICs) developed on natural biopolymer scaffold, poly(β-L-malic acid), with covalently attached a-CTLA-4 and/or a-PD-1 for delivery across the BBB and activation of local brain anti-tumor immune response in glioma-bearing mice. NIC treatment of mice bearing intracranial GL261 glioblastoma (GBM) resulted in an increase of CD8+ T-cells with a decrease of T regulatory cells (Tregs) in the brain tumor area. Survival of GBM-bearing mice treated with combination of NICs was significantly longer compared to animals treated by single checkpoint inhibitor-bearing NICs or free a-CTLA-4 and a-PD-1. Our study demonstrates trans-BBB delivery of nanopolymer-conjugated checkpoint inhibitors as an effective treatment of GBM via activation of both systemic and local brain tumor immune response.

---

Glioblastoma multiforme (GBM), the most aggressive brain glioma, has seen only small treatment advances in the last 30 years. Treatment options are limited, in part because of inefficient drug delivery across the blood-brain barrier (BBB). Immunotherapy is one of the fastest developing approaches in clinical oncology. However, the unique immune enviroment of the central nervous system (CNS) needs consideration when pursuing immunotherapeutic approaches for gliomas. Advances in nanotechnology allow the design of nanocarriers for targeted delivery of cancer vaccines and chemo-immunotherapies^1^. We combine achivements in nanotechnology and immunotherapy into a new approach to deliver drugs across the BBB and treat glioblastoma multiforme.

Blockade of cytotoxic T-lymphocyte-associated antigen 4 (CTLA-4) using the antagonistic monoclonal antibody (mAb) ipilimumab was the first strategy to achieve a significant clinical benefit for stage IV melanoma patients^2, 3^ Humanized mAbs against immune system response modulators CTLA-4 (ipilimumab), and programmed cell death-1 (PD-1) (pembrolizumab and nivolumab), received FDA approval. Their effect is due to the suppression of T regulatory cells (Tregs) and activation anti-tumor immune response by cytotoxic T lymphocytes (CTLs). Systemic administration of CTLA-4 or PD-1 and programmed cell death ligand 1 (PD-L1) mAbs can suppress some tumors, but has low efficacy against brain and breast tumors^4, 5^

Despite growing evidence to support an interaction between the CNS and the systemic immune system^6^, clinical trials using nivolumab and ipilimumab in GBM showed serious safety issues, but not a significant anti-tumor treatment effect^7^. Although CTLA-4 and PD-1 antibodies do not cross the BBB^8, 9, 10^, a modest efficacy has been demonstrated against GBM, possibly due to general immune system activation. A recently published review^11^ summarized the results of several immunotherapy clinical trials in glioma: 28 clinical trials for vaccines (e.g., a peptide vaccine that targets EGFRvIII or IDH1); 13 clinical trials completed for oncolytic viruses; 15 phase III clinical trials for checkpoint inhibitors (e.g., CheckMate 143 trial) and genetically modified T cells expressing chimeric antigen receptors (CAR-T cells). Unfortunately, no treatment so far has been superior to the standard-of-care for GBM, represented by temozolomide/radiation therapy with 14.6 months survival. This demonstrates serious unmet need for a novel technological approach. Here we use a versartile drug carrier, poly(β-L-malic acid) (PMLA), a natural polymer obtained from the slime mold *Physarum polycephalum*^12^, to deliver covalently conjugated CTLA-4 and PD-1 antibodies (a-CTLA-4 and a-PD-1) to brain tumor cells. PMLA-based nanotherapeutics can target brain tumors by crossing the BBB using transferrin receptor (TfR) mediated transcytosis^12, 13^. To our knowledge, this is the first successful use of polymer-based nanocarriers with covalently attached immunotherapeutics to activate local immune response and treat brain tumors.

## Results

### NICs synthesis

We covalently attached checkpoint inhibitor antibodies, a-CTLA-4 IgG2b or a-PD-1 IgG, to the PMLA backbone to reach stability for plasma circulation. mPEG5000 was attached for solubility and stability, anti-mouse TfR antibody (a-msTfR), to cross the BBB, and trileucine (LLL), for stabilization of PMLA against hydrolytic degradation^14^, hydrophobization, and for endosomolytic drug delivery^15, 16^. Synthesis has been adapted from previous studies^12, 13, 14, 15, 16^ with new analysis specific to our NICs (Fig. 1) that have met physicochemical and immunological criteria for translational usage.

**Figure 1.**
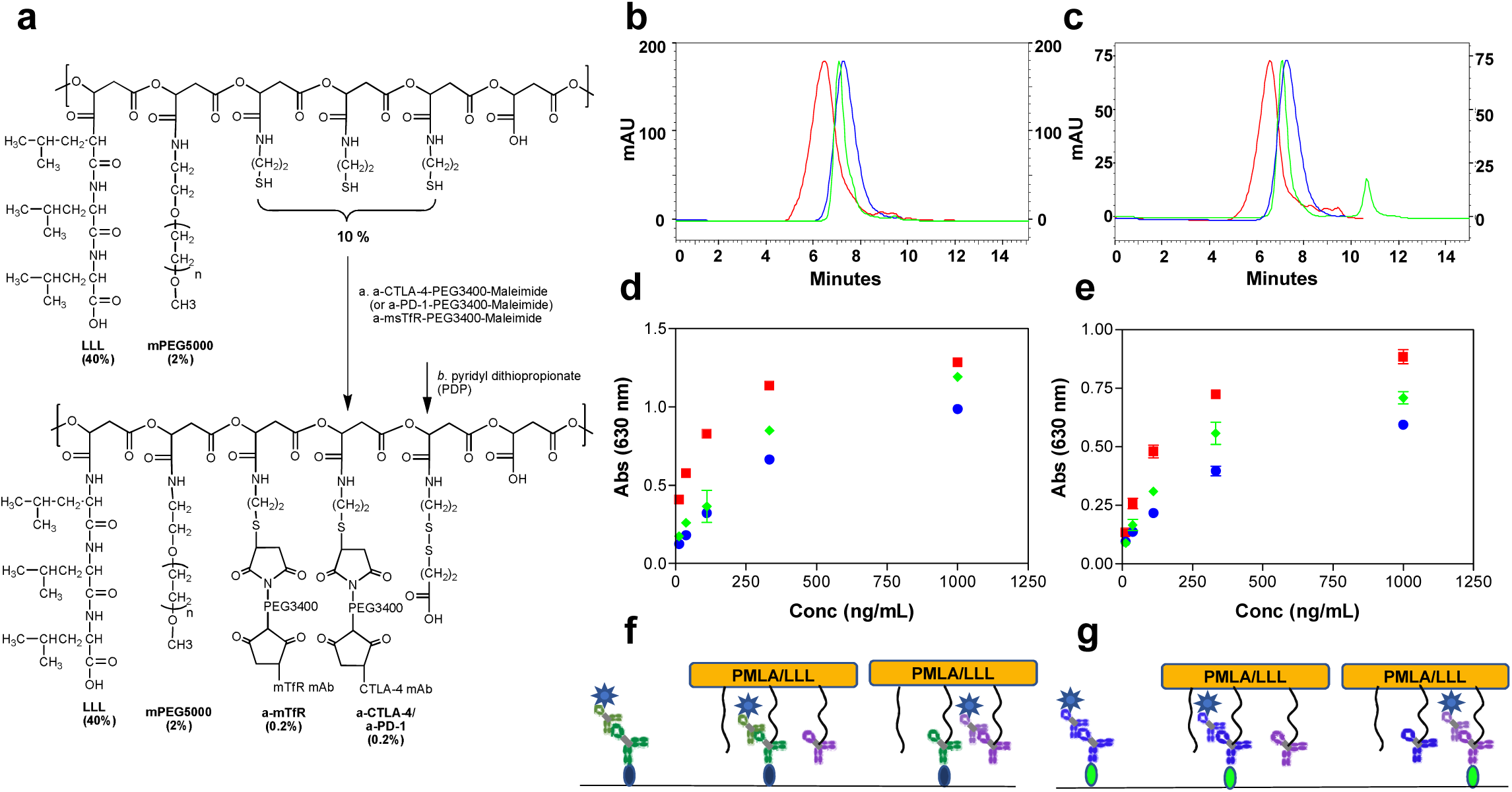
Synthesis and characterization of NICs. **a,** Synthesis of P/mPEG5000(2%)/LLL(40%)/a-msTfR(0.2%)/a-CTLA-4 (0.2%). The upper structure represents the pre-conjugate and the lower structure represents the final nanoconjugate with antibodies and 3-(2-pyridyldithio) propionate (PDP). **b,** SEC-HPLC analysis of P/mPEG5000(2%)/LLL(40%)/a-msTfR(0.2%)/a-CTLA-4(0.2%). Right blue peak, the pre-conjugate P/mPEG5000(2%)/LLL(40%)/MEA(2%); middle (green) peak, a-CTLA-4; red peak, the synthesized P/mPEG5000(2%)/LLL(40%)/a-msTfR(0.2%)/a-CTLA-4(0.2%) **c,** SEC-HPLC analysis of P/mPEG5000(2%)/LLL(40%)/a-msTfR(0.2%)/a-PD-1(0.2%). Right blue peak, the pre-conjugate P/mPEG5000(2%)/LLL(40%)/MEA(2%); middle (green) peak, a-PD-1; red peak, the synthesized P/mPEG5000(2%)/LLL(40%)/a-msTfR(0.2%)/a-PD-1(0.2%). **d-e,** Validation of simultaneous conjugation and activity of a-msTfR and a-CTLA-4/a-PD1 on a single platform by ELISA of P/aCTLA-4 (**d**) and P/a-PD-1 (**e**). a-CTLA-4 and a-PD-1 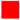, a-CTLA-4 and a-PD-1 on conjugate
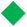
a-msTfR
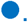
**f,** Illustration of ELISA method used in **d**. **g,** illustration of ELISA method used in **e**.

Synthesis of NICs for polymer conjugated a-CTLA-4, as an example, involved synthesis of pre-conjugate P/mPEG5000(2%)/LLL(40%)/MEA(10%), chemical activation of mAb maleimide, a-CTLA-4-PEG3400-maleimide, and a-msTfR-PEG3400-maleimide, and conjugation of pre-conjugate with activated mAbs through thiol ether formation (Fig. 1a). This was followed by blocking residual free thiol groups. Size-exclusion HPLC (SE-HPLC) confirmed conjugation (Fig. 1b-c). Fourier transform infrared spectroscopy (FTIR) and pull-down ELISA further validated the structure of P/mPEG5000(2%)/LLL(40%)/a-msTfR(0.2%)/a-CTLA-4(0.2%) (P/a-CTLA-4) (Fig. 1d, f; Supplementary Fig. 1a, c) and P/mPEG5000(2%)/LLL(40%)/a-msTfR(0.2%)/a-PD-1(0.2%) (P/a-PD-1) (Fig. 1e, g; Supplementary Fig. 1b, d).

Free a-CTLA-4, a-PD-1, and a-msTfR showed somewhat higher binding affinity toward their respective antigens on a plate surface compared with polymer-bound antibodies, due to the bulkier size of the NICs. In addition, we proved the presence of pairs of two antibodies (i.e. a-CTLA-4 and a-msTfR or a-PD-1 and a-msTfR) within one single polymer chain using pull-down ELISA (Fig. 1f-g, Supplementary Fig. 1c-d). The ELISA intensity of a-msTfR on P/a-CTLA-4 conjugate was comparable to that of a-CTLA-4 on the same conjugate, confirming the presence of both a-msTfR and a-CTLA-4 on the conjugate (Fig. 1d). Similar results were obtained for P/a-PD-1, when the surface was coated with PD-1 (Fig. 1f), and for P/a-CTLA-4 and P/a-PD-1, when the surface was coated with msTfR (Supplementary Fig. 1a-b). ELISA results confirmed the reactivity of each antibody and the presence of each antibody in the conjugates. FTIR analysis for P/a-CTLA-4 shows peaks at the O-H stretching frequencies of 2880 cm^-1^ (carboxylic acid O-H) that can be seen in both pre- and final nanoconjugate, as well as peaks that are present at the a-CTLA-4 spectrum, which are also present in the final NIC (mainly peaks at 3270 and 2953 cm^-1^, which are N-H and O-H stretching frequencies, and lower frequencies peaks at 1120 and 1630 cm^-1^) (Supplementary Fig. 2a). A similar trend could also be seen for P/a-PD-1, with significant specific peaks at 2875, 1664, 1031, and 942 cm^-1^ (Supplementary Fig. 2b). The analysis of the FTIR spectra suggested that the antibodies were conjugated successfully to the pre-conjugate.

ζ-Potentials of NICs were in the range of -9.9 to -11.0 mV, reflecting the design and the intrinsic antibody charges, hydrodynamic size was in the range of 28.0 to 28.5 nm by intensity, in agreement with design and molecular masses of constituents and the absence of aggregates, and endotoxin level was reduced below 0.1 EU/mL by phase separation method^17^ (Tab. 1).

### NICs cross the BBB and reach the tumor interstitium

Rhodamine-labeled NICs (Fig. 2a-b) were detected in the brain of tumor-bearing mice, 4h after intravenous (i.v.) injection. NICs were distributed throughout the tumor area, mostly outside the blood vessels, with only occasional presence in the vessels. Rhodamine-labeled a-CTLA-4 and a-PD-1 could not be detected in the brain at 4h and 6h after i.v. injection, indicating their inability to cross the BBB and reach the brain parenchyma (Fig. 2c-d). Conversely, NICs cross the BBB, hence allowing conjugated mAbs to bind to CTLA-4 and PD-1 and modulate the immune response in the tumor area (Fig. 2e-f).

**Figure 2.**
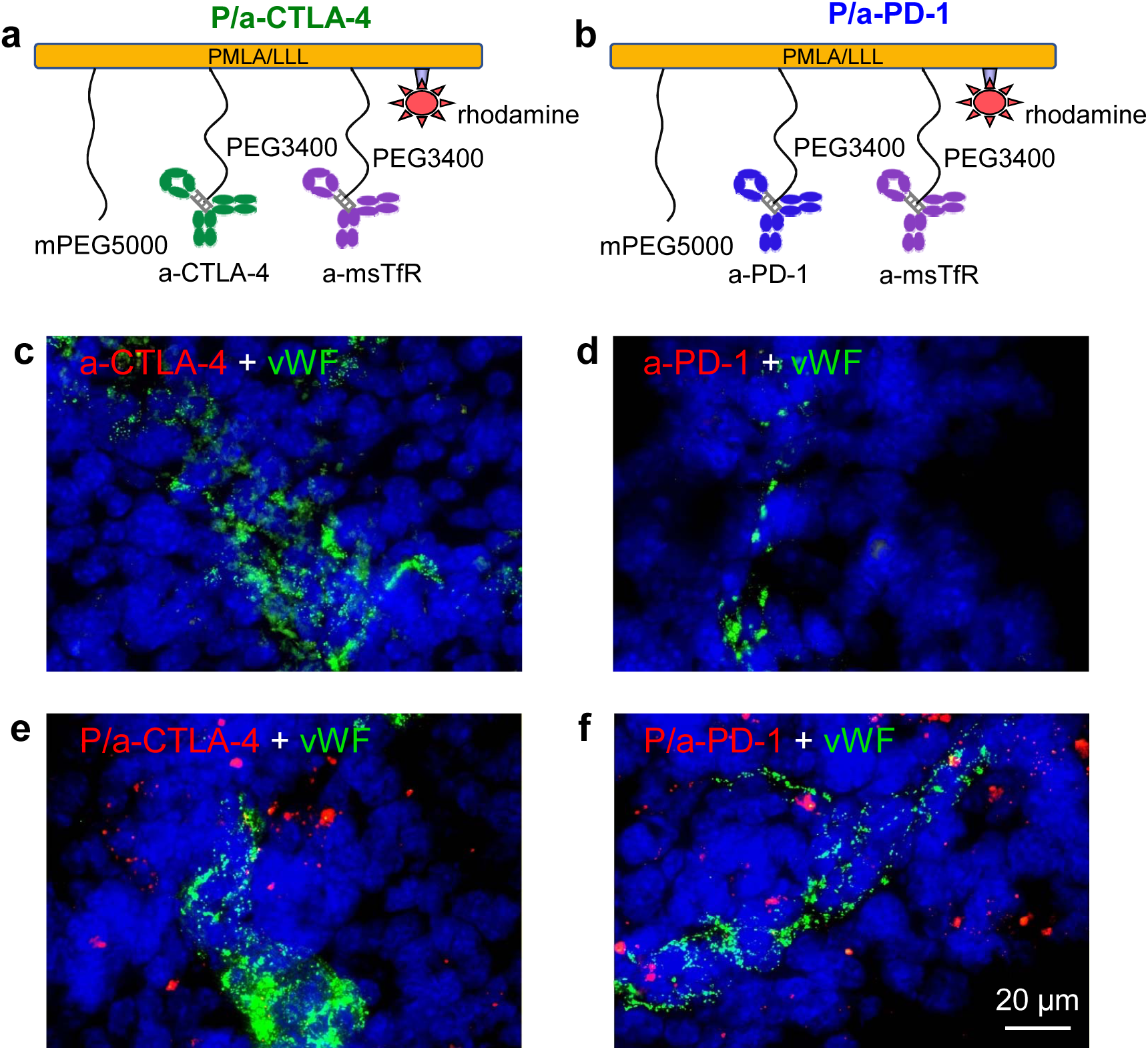
Fluorescently labeled NICs cross the BBB into intracranial glioma. **a,** Structure of a nanoconjugate containing the platform PMLA/LLL/mPEG5000(2%)/rhodamine(1%), PEG3400/a-msTfR (0.2%), and PEG3400/a-CTLA-4 (0.2%). **b,** NIC with the same platform composition as in A containing PEG3400/a-msTfR (0.2%), and PEG3400/a-PD-1(0.2%). **c-d,** Free (non-conjugated) rhodamine-labeled antibodies (a-CTLA-4 and a-PD-1) are not detectable in the brain. **e-f,** NICs containing a-CTLA-4 or a-PD1 (red) are distributed outside the blood vessels in the tumor parenchyma. Blood vessels are stained with anti-von Willebrand factor (vWF) antibody (green). Scale bar on bottom right panel applies to all other images.

### NIC treatments increase survival of GL261 glioblastoma-bearing mice

Most preclinical studies with checkpoint inhibitors used intraperitoneal (i.p.) administration of therapeutics^18, 19^, to avoid anaphylactic shock after i.v. injection. However, systemic i.v. drug administration is widely considered as the clinically accepted method in brain cancer treatment. In animal models, we observed a rapid and fatal hypersensitivity reaction with repeated i.v. injections of a-CTLA-4, a-PD-1, and NICs during our initial experiments. Therefore, to avoid acute immune-mediated anaphylaxis-like side effects, premedication must be used^20, 21,22^.

Although the first two i.v. treatments did not cause any noticeable side effects, mice experienced a severe drop in body temperature, piloerection, loss of spontaneous activity, dyspnea, and lethargy 15-20 min after the following injections. As much as 20% of mice died after the third treatment and up to 100% after the fifth treatment. Different regimens were explored to reduce treatment dosage and frequency, however, without eliminating toxicity (data not shown). To counteract these side effects, 10 mg/kg antihistamine Triprolidine and/or 5 mg/kg platelet activating factor (PAF) antagonist CV6209 were administered prior to each drug injection following the first one. Although both Triprolidine and CV6209 reduced side effects to some level, only when administered in our new combination regimen, they prevented side effects in 100% of cases (data not shown). Premedication allowed 5 repeated i.v. treatments using a 10 mg/kg antibody dose, fully comparable to the current clinical dosage of checkpoint inhibitor mAbs.

Survival of mice bearing intracranial GBM GL261 and treated with free mAbs or NICs alone or in combination was investigated. Both a-CTLA-4 and a-PD-1 failed to increase survival compared to PBS treatment. Crucially, both P/a-CTLA-4 and P/a-PD-1 (Fig. 3a-b) significantly improved animal survival compared to PBS (p=0.0076 and p=0.0017, respectively) or the respective free antibody treatment (p=0.0362 and p=0.0036, respectively) (Fig. 3d-e). A combination of P/a-CTLA-4+P/a-PD-1 further improved survival in GL261 tumor-bearing mice compared to PBS (p<0.0001), a-CTLA-4 (p<0.0001), a-PD-1 (p<0.0001), P/a-CTLA-4 (p<0.0001), and P/a-PD-1 (p=0.0033) (Fig. 3d-e; Supplementary Fig. 3; Supplementary Video).

**Figure 3.**
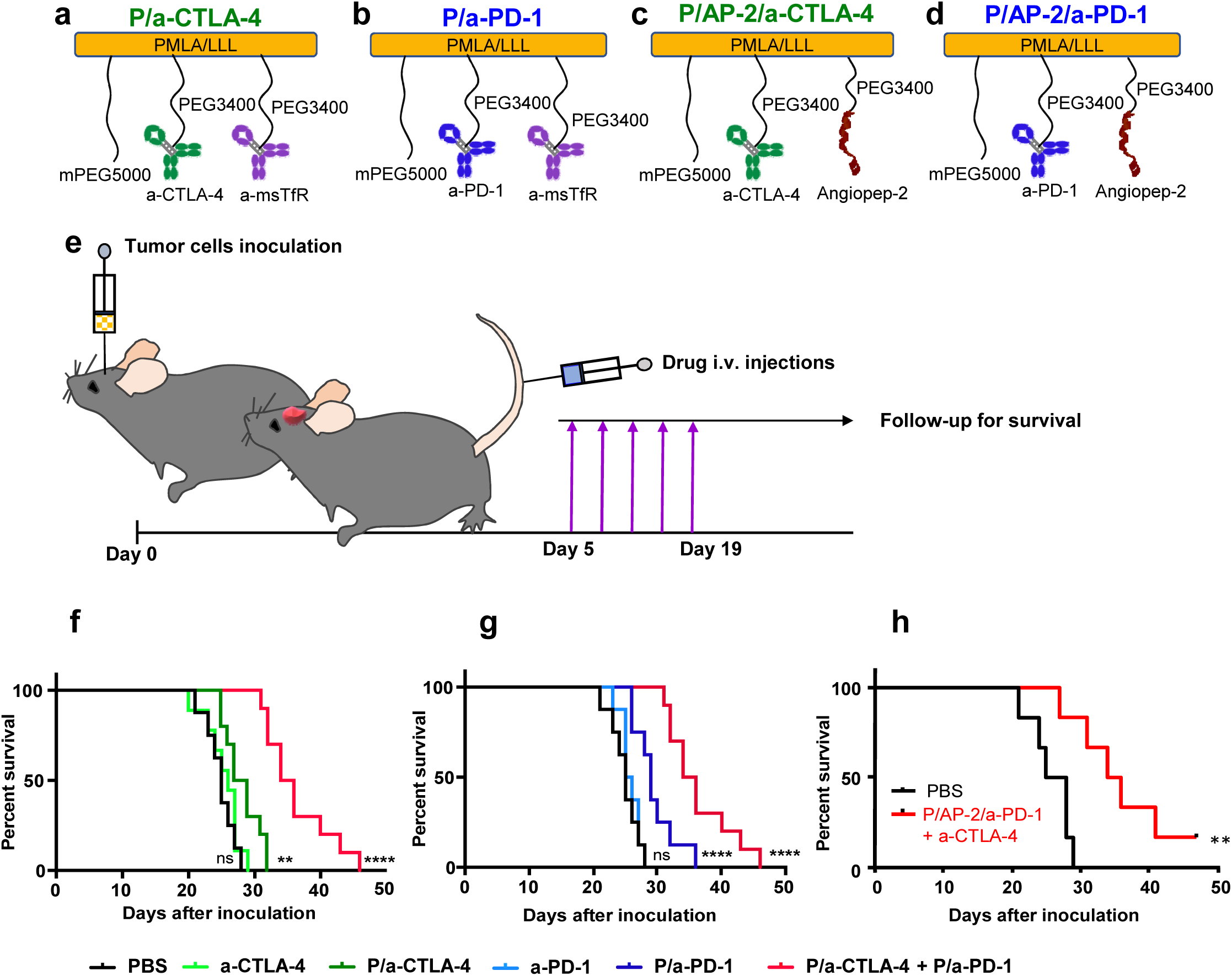
Treatment of mice bearing intracranial GL261 glioblastoma tumors with NICs. **a,** Structure of treatment nanoconjugate containing PMLA/LLL as a backbone, 2% mPEG5000, a-msTfR and a-CTLA-4. **b,** Structure of treatment nanoconjugate containing PMLA/LLL as a backbone, 2% mPEG5000, a-msTfR and a-PD-1. **c,** Schematic depicting experiment workflow: tumor cells are inoculated intracranially; five i.v. drug treatments are started 5 days after inoculation after which animals are followed for survival. **d,** Kaplan-Meier plot of animal survival after treatment with PBS, a-CTLA-4, P/a-CTLA-4, and P/a-CTLA-4+P/a-PD-1. Treatment with P/a-CTLA-4 and P/a-CTLA-4+P/a-PD-1 significantly increased survival compared with free antibody and PBS. **e,** Kaplan-Meier survival curve for animals treated with PBS, a-PD-1, P/a-PD-1, and P/a-CTLA-4+P/a-PD-1. Both NIC treatments significantly improved survival compared to free antibody and PBS. P-values were obtained using a log-rank (Mantel-Cox) test where ns = non-significant, ** = p<0.01, **** = p<0.0001.

### NICs activate local brain Immune response

Flow cytometry analysis showed an expansion of the CD8+ T cell population in the tumor tissue after treatment with NICs, especially in animals treated with P/a-PD-1, compared to PBS and free mAb treatments (Fig. 4a). On the contrary, the Treg fraction (CD4+FoxP3+) decreased in all NIC groups compared to PBS and free mAb (Fig. 4b). CD8+ T cells are cytotoxic lymphocytes that carry out the attack and kill tumor cells via different pathways, whereas Tregs are regulatory lymphocytes modulating the immune system to maintain tolerance to self-antigens. Animals with survival longer than the median survival for their treatment group showed larger differences in Treg population between treatments (Fig. 4c). Combination therapy achieved the highest reduction in the Treg fraction (Treg fraction = 32.87% for P/a-CTLA-4 + P/a-PD-1 *vs.* 51.43% for PBS), which is in line with significantly increased animal survival.

**Figure 4.**
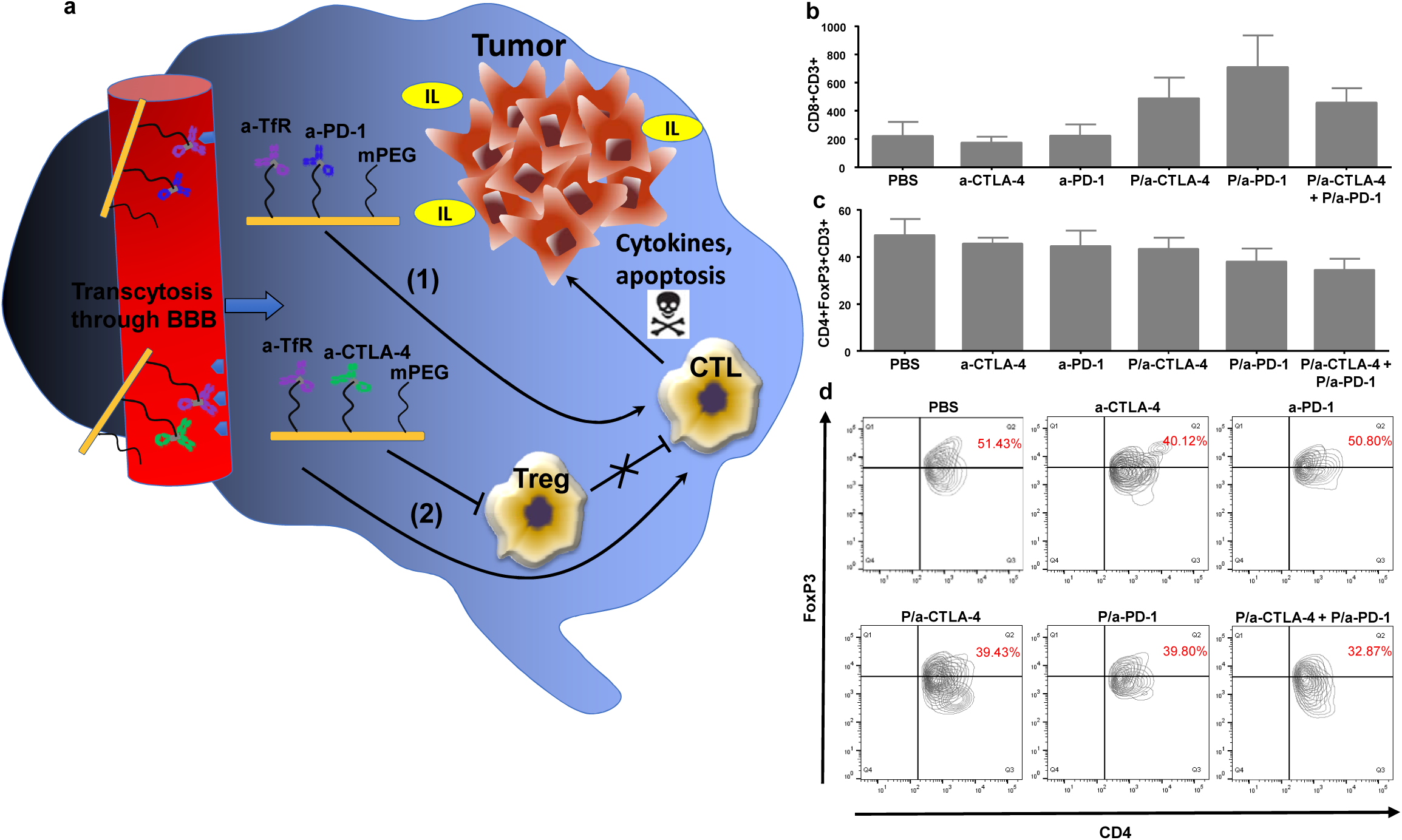
Effect of NICs on CTL and Treg populations in GL261 intracranial tumors. **a,** Nano immunology concept for treatment. PD-1 blockade leads to CTL activation (1). Blocking CTLA-4 pathway prevents CTL inhibition (2) **b,** Flow cytometry analysis showing increase in CD8+ T cells in NIC treated groups compared to PBS and free antibodies. **c,** Flow cytometry analysis displaying a decrease in CD4+FoxP3+ T cells after treatment with NICs compared with free antibody and PBS treated animals. **d,** Flow cytometry analysis of long survivor animals (survival longer than median survival for the relative group) showing a Treg fraction (shown in Q2) decrease in animals treated with NICs compared with those treated with PBS and free a-CTLA-4 or a-PD-1. N=4-10 in each group.

To independently confirm flow cytometry data, the local immune system response was investigated with specific CD8 and CD4+FoxP3 immunofluorescence staining of cryosectioned brain tumor tissues. For CD8 staining, we used FIJI software to analyze 4032 cells in tumors from 12 mice representing different treatment groups (Fig. 5a-g). On average, each analyzed image contained 106 ± 22 cells, and cell densities were uniform for all data included in our analysis. We observed a significant increase in the incidence of CD8+ T cells in tumor tissue (F=5.826; p=0.0006) (Fig. 5g). In particular, the percentage of CD8+ T cells in drug *vs.* PBS groups was significantly increased following treatments with P/a-PD-1 (p=0.0213) and P/a-CTLA-4 + P/a-PD-1 (p=0.0005). No other treatment showed a statistically significant change in the incidence of the CD8+ T cells when compared to the PBS. However, a statistically significant increase in the percentage of CD8+ T cells was also observed when comparing a-CTLA-4 (p=0.0118) and P/a-CTLA4 (p=0.0340) with the P/a-CTLA-4 + P/a-PD-1 condition (Fig. 5g). This result confirms that the NIC combination treatment is the most effective at recruiting CD8+ T cells into the tumor tissue.

**Figure 5.**
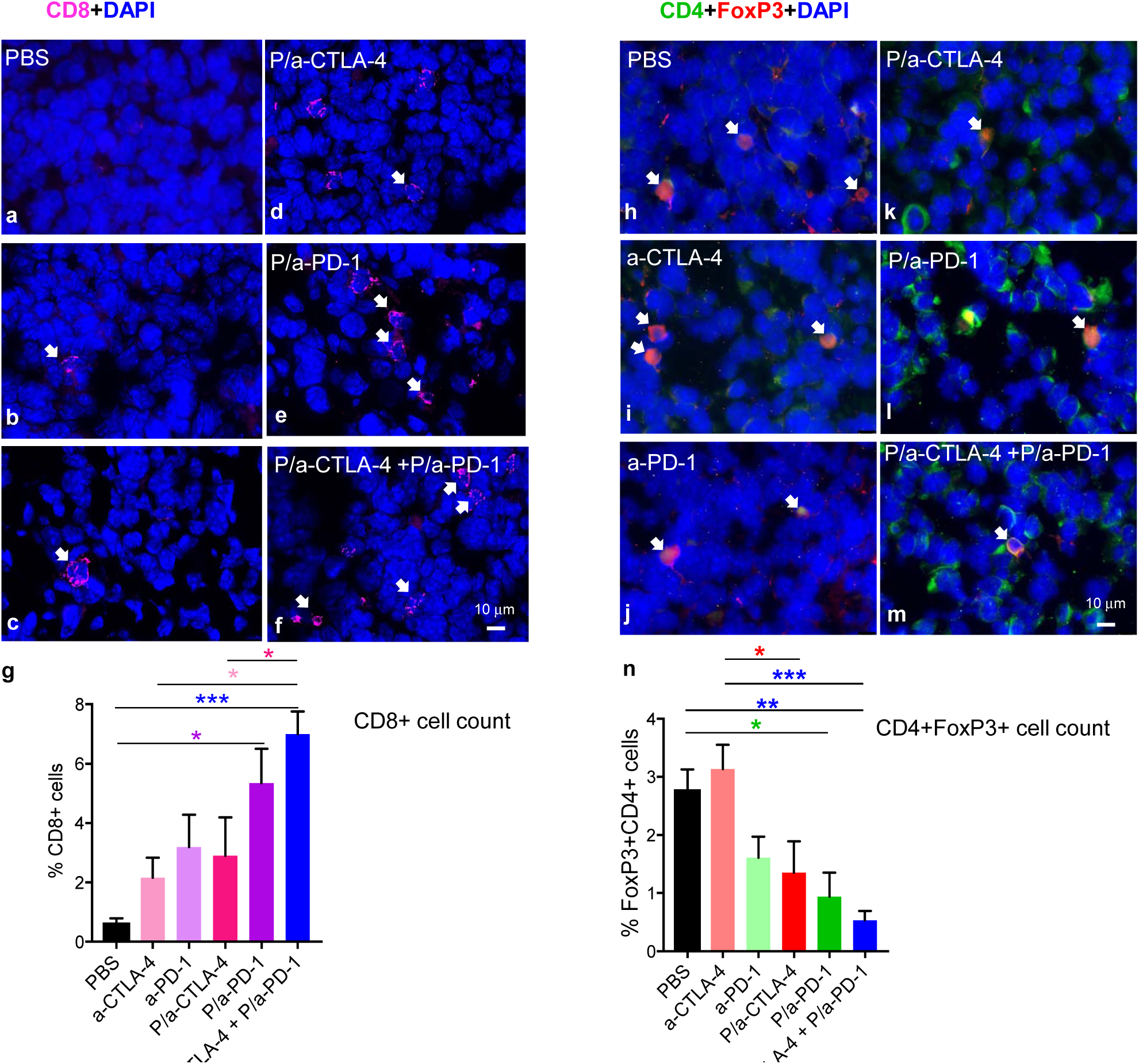
Increase of CD8+ and decrease of CD4+FoxP3+ T cell populations in GL261 intra-cranial tumors after NIC treatment. Immunostaining of CD8+ T cells in tumors treated with **a,** PBS, **b,** a-CTLA-4, **c,** a-PD-1, **d,** P/a-CTLA-4, **e,** P/a-PD-1, and **f,** P/a-CTLA-4+P/a-PD-1. Arrows indicate CD8+ lymphocytes (magenta). Blue indicates nuclear staining with DAPI. Images are shown at 63X magnification. Scale bar on bottom right panel applies to all other images. **g,** Treatments with P/a-PD-1 and P/a-CTLA-4+P/a-PD-1 result in a statistically significant increase of CD8+ lymphocytes inside the tumor. * = p<0.05; *** = p<0.001 (one-way ANOVA + Tukey’s post-hoc test). Immunostaining of CD4+FoxP3+ T cells in tumors treated with **h,** PBS, **i,** a-CTLA-4, **j,** a-PD-1, **k,** P/a-CTLA-4, **l,** P/a-PD-1, and **m,** P/a-CTLA-4+P/a-PD-1. Arrows indicate CD4+FoxP3+ lymphocytes (yellow). CD4+ staining is in green, FoxP3+ staining is in red, DAPI nuclear staining is in blue. Images are shown at 63X magnification. Scale bar in bottom right panel applies to all other images. **n,** P/a-PD-1 and P/a-CTLA-4+P/a-PD-1 treatments significantly decrease the number of CD4+FoxP3+ lymphocytes in the tumor. * = p<0.05; ** = p<0.01; *** = p<0.001 (oneway ANOVA + Tukey’s post-hoc test).

To determine the effect of our drug treatments on CD4+FoxP3+ cells in tumor tissue, we measured the number of CD4/FoxP3 double immunolabeled T cells. Cells that were exclusively FoxP3+ were not included in our analysis. We analyzed 4736 cells in tumor tissues from 12 mice (2 mice per treatment group) (Fig. 5h-n). On average, each analyzed image contained 118 ± 20 cells; cell densities were uniform. We observed a significant decrease in the incidence of CD4+FoxP3+ T cells in tumor tissue (F=7.02; p=0.0001) (Fig. 5n). Specifically, we observed a significantly lower percentage of CD4+FoxP3+ T cells following treatment with P/a-PD-1 (p=0.0235) and P/a-CTLA-4 + P/a-PD-1 (p=0.0013) when compared to PBS (Fig. 5n). Strikingly, this result is almost exactly inverse to our observation related to CD8+ T cells. No other drug treatment showed a statistically significant change in the incidence of the CD4+FoxP3+ T cells when compared to PBS. However, a larger reduction of CD4+FoxP3+ T cells was again observed when comparing the a-CTLA-4 with the P/a-CTLA-4 (p=0.0311) and P/a-CTLA-4 + P/a-PD-1 (p=0.0006) treatments (Fig. 5n). These data again suggest that our combination treatment is the most effective at inhibiting CD4+FoxP3+ T cells in the tumor tissue.

Collectively, flow cytometry and histology results confirmed that the treatment with NICs stimulated the local brain immune system allowing cytotoxic T cells to attack the tumor.

### NICs increase systemic immune response

When activated, CD4+ and CD8+ T cells produce inflammatory cytokines. To confirm systemic immune activation following NIC treatment, serum cytokine levels were measured by multiplex assay. Treatment with P/a-CTLA-4 showed increase of serum levels for multiple interleukins: IL-1β (p<0.01), IL-2 (p<0.01), IL-4 (p<0.01), IL-5 (p<0.01), and IL-12(p70) (p<0.01) (Supplementary Fig. 3a-d, g). In contrast, after treatment with P/a-PD-1, only serum levels of IL-2 increased significantly (p<0.01) in comparison with PBS-treated group (Supplementary Fig. 3b). The highest increase for all 9 cytokines measured was seen for the combination treatment with P/a-CTLA-4 + P/a-PD-1 vs. PBS treatment: IL-1β (p<0.0001), IL-2 (p<0.0001), IL-4 (p<0.001), IL-5 (p<0.01), IL-6 (p<0.01), IL-10 (p<0.01), IL-12(p70) (p<0.05), IFN-γ (p<0.01), and TNF-α (p<0.05) (Supplementary Fig. 3a-i). These data suggest that the systemic immune response was also activated by our drug treatment.

## Discussion

The Immuno-Oncology Translational Network is the number one program on the list of multi-disciplinary, collaborative approaches to accelerate understanding of the tumor immune microenvironment in response to NIH/NCI Blue Ribbon Panel report^23^.

We developed polymeric NICs with covalently attached checkpoint inhibitor mAbs a-CTLA-4 and a-PD-1. They were able to cross the BBB via the proven delivery system using polymer-conjugated a-msTfR^15, 16^, and elicit anti-tumor immune responses in brain glioma in a mouse model. NICs were immunochemically investigated for their biological ability to bind substrates, mimicking the natural conditions for naked a-CTLA-4 and a-PD-1. The analytical results were satisfactory to move to the delivery and therapeutic steps.

The glioma immune microenvironment is very complex, but, with proper checkpoint inhibitor delivery through the BBB, the basic concept for immune treatment still holds true: disrupting the PD-1/PD-L1 and CTLA-4/B7-1 complex formation in the tumor is key to the improved survival of glioma-bearing mice.

Checkpoint inhibitors have shown clinical efficacy for the treatment of “hot” immunogenic tumors, such as melanoma and other cancers with high rate of lymphocyte infiltration, when GBMs are considered “cold” and poorly infiltrated with lymphocytes^11,24^. The most significant obstacles in the treatment with checkpoint inhibitors are tumor resistance and toxicity^25^, as well as their inability to cross biological barriers such as the BBB^8^. GBM, a very aggressive tumor with short survival and with limited and poorly effective treatment options, presents a particular challenge for drug delivery because of its location in the CNS and the necessity for the drugs to cross the BBB. A recent study has described survival improvement and immune system activation in glioma-bearing mice treated with free checkpoint inhibitor mAbs (a-PD1 and a-CTLA-4)^26^. Although such antibodies do not bind known receptors capable of BBB transcytosis and are unlikely to function effectively in the brain, they may still provide a modest activation of the brain immune system, possibly working through systemic immune stimulation^8, 10, 25^

For the first time, we demonstrated the efficacy of NICs carrying a-PD-1 and a-CTLA-4 in treating GBM in a murine model vs. free a-CTLA-4 and a-PD-1. We established that NICs, as single nano agent therapies and, especially, as a combination of P/a-PD-1 + P/a-CTLA-4, cross the BBB and significantly increase survival of GL261 tumor-bearing mice activating the brain resident immune system compared to treatment with free antibodies (Supplementary Video).

The activation of the resident immune system following NIC treatments was evidenced by a local increase in CD8+ T cells, which are cytotoxic effector T cells that attack the tumor, and a reduction of Tregs, which suppress activation and proliferation of effector T cells. Tregs population in the brain tumor significantly decreased after NIC treatments compared to PBS and free mAbs.

As a consequence of immune system activation, serum levels of IL-1β, IL-2, IL-4, IL-5, IL-6, IL-10, and IFN-γ were significantly increased after combination treatment with P/CTLA-4 + P/PD-1 compared to other treatments. Cytokines are produced by T cells to regulate immune response and to kill cancer cells through apoptotic and non-apoptotic pathways^27^.

In general, IL-1β, IL-2, IL-12, IFN-γ, and TNF-α are part of T helper 1 (Th1) response, whereas IL-4, IL-5, IL-6, and IL-10 are part of T helper 2 (Th2) response. IL-2 secretion promotes T and B lymphocyte activity, enhances anti-tumor immunity, stimulates microglia, and regulates Tregs^28, 29, 30^ IFN-γ is mainly produced by CD4+ and CD8+ T cells, NK cells, and microglia to boost the cytotoxic immune response^30, 31^. In a self-stimulating loop, the microglia can also secrete IL-12, activating NK and stimulating T cells^32^. Conversely, IL-10 is a multifunctional immune cytokine with anti-angiogenic properties^33, 34^ Finally, cytokines like IL-1β and TNF-α can suppress tumors via stimulation of cell-mediated humoral immune reaction^35^. In our study, all these cytokines are elevated in serum of animals treated with NIC combination, supporting increased cytotoxic activity of the immune system. Some of these cytokines have been used in anti-cancer treatment: IL-4 for GBM^36, 37^ IL-12 for breast cancer brain metastasis^38^, IL-2 for melanoma brain metastasis^39, 40^, and IFN-α and IL-2 for renal cell carcinoma^41^. CTLA-4 and PD-1 suppression also increased IL-4, IL-5, IL-6, and IL-10, which is a known effect of such treatments in other cancer types^42, 43, 44^.

Two pathways of anaphylaxis are known in mice: one is caused by antigen crosslinking of IgE on mast cells followed by release of histamine^45, 46^, and the second one is mediated by IgG1 and basophil or neutrophil release of PAF^20, 47^. Immunotherapy with a-CTLA-4 and a-PD-1 caused anaphylaxis-like side effects after repeated administrations. However, we observed absence of adverse effects after premedication consisting of anti-histamine, Triprolidine, and PAF antagonist, CV6209, previously shown to prevent anaphylaxis in other contexts^22^.

In our study, premedication safely allowed for 5 repeated i.v. administrations at a therapeutic dosage comparable with the clinical settings. PAF inhibitor drugs are currently available for clinical use in cardiac rehabilitation and could be adopted in future clinical trials to mitigate adverse immune-mediated events related to checkpoint inhibitor therapy^22^.

Overall, our results show that BBB-crossing NICs stimulate the brain resident immune system, prompting the proliferation of CD8+ T-cells and triggering the release of several cytokines, thus orchestrating immune response against glioblastoma. These NICs provide a novel and useful tool for the delivery of immunotherapy and targeted therapies to brain tumors. They may be used for treatment of primary brain tumors and brain metastases, where current treatment is very inefficient^48, 49^ due to the inability to deliver therapeutics through the BBB and activate brain privileged immune system.

A new regimen of premedication appears to alleviate adverse immune-mediated effects upon repeated i.v. injections of NICs, allowing the use of this clinically relevant route of administration in animal studies of checkpoint inhibitors.

## Methods

### Reagents

Polymalic acid (PMLA) with molecular mass 50,000 D (SEC-HPLC/polystyrene sulfonate standards, polydispersity 1.2) was isolated from the culture supernatant of *Physarum polycephalum* M3CVII as previously described^14, 50^. Mal-PEG3400-Mal and mPEG5000-NH2 were obtained from Laysan Bio (Arab, AL). Cy5.5 maleimide was obtained from Lumiprobe (Hallandale Beach, FL, USA). Superdex G-75 was obtained from GE Healthcare (Uppsala, Sweden). InVivoMAb anti-mouse PD-1 (clone j43, Isotype Armenian hamster IgG) was from BioXcell (West Lebanon, NH) and mouse anti-mouse a-CTLA-4 IgG2b (clone 9D9) was from Bristol-Myers Squibb (Redwood City, CA).

### Synthesis of NICs

#### Synthesis of P/mPEG5000(2%)/LLL(40%)/2-mercapto-ethylamine (pre-conjugate)

Pre-conjugate P/mPEG5000(2%)/LLL(40%)/2-mercapto-ethylamine was synthesized based on the method previously described^12, 13, 15^. At first, the pendant polymer carboxylates were converted into highly reactive NHS ester from N-hydroxysuccinimide (NHS), dicyclohexylcarbodiimide (DCC) (1:1:1 molar ratio for PMLA-COOH:NHS:DCC) in acetone/DMF (1:1, 20°C, 2 h). The ester was then substituted by amidations following the addition of mPEG5000-NH2 (2% equivalent to the total malyl groups in 200 μL DMF; 1 h), H-Leu-Leu-Leu-OH (LLL) (40% dissolved in DMF with 1.1 equivalent of trifluoroacetic acid to LLL for solubilization, added in 6 portions at 10min intervals, each followed by aliquots of 1.1 equivalent of trimethylamine to LLL), 2-mercaptoethylamine (MEA) (10%, DMF, 30 min, together with DTT 2 equivalent to MEA, followed by 1 equivalent triethylamine). The completion of each reaction was controlled by ninhydrin test. After hydrolysis of unreacted NHS ester by the addition of phosphate buffer (100 mM, pH 6.8), precipitation of dicyclohexylurea and removal by filtration, the pre-conjugate P/mPEG5000(2%)/LLL(40%)/MEA(10%) was purified over PD-10 column (GE Healthcare, Pasadena, CA) and lyophilized (Fig. 1a).

#### Synthesis of a-CTLA-4-PEG3400-Maleimide, a-PD-1-PEG3400-Maleimide and ams TfR-PEG3400-Maleimide

Anti-mouse TfR mAb (a-msTfR; 10 mg), anti-mouse CTLA-4 mAb (a-CTLA-4; 10 mg) and anti-mouse PD-1 mAb (a-PD-1; 10 mg) were reduced with Tris (2-carboxy ethyl) phosphine hydrochloride (TCEP) (5 mM, 30 min, 20°C). After removal of excess TCEP on PD-10 column (GE Healthcare), the reduced antibody was conjugated with Mal-PEG3400-Mal (10 mmole, EDTA 5 mM, 1 mL PBS, 30 min). Successful conjugation was confirmed by size exclusion HPLC (SE-HPLC). The maleimide derivatives of antibodies were concentrated by centrifuging filtration with Vivaspin 20 (cutoff 30 kD, Sartorius Stedim Biotech, Concord, CA), and the product was purified over Sephadex G75 with phosphate buffer 100 mM pH 6.3 as eluent.

#### Synthesis of P/mPEG5000(2%)/LLL(40%)/a-CTLA-4(0.2%)/a-msTfR(0.2%) and P/mPEG5000(2%)/LLL(40%)/a-PD-1(0.2%)/a-ms TfR(0.2%)

To the mixture of a-CTLA-4-PEG3400-Maleimide (or a-PD-1-PEG3400-Maleimide) and a-msTfR-PEG3400-Maleimide (10 mg each, equal to 0.2% equivalent to total malyl groups) in 4 mL 100 mM sodium phosphate buffer (100 mM, pH 6.3) pre-conjugate (P/mPEG5000(2%)/LLL(40%)/2-mercapto-ethylamine 2.5 mg/mL; 11.7 mg, in 4 mL of phosphate buffer) was added dropwise. Successful conjugation (30 min, 20°C) was confirmed by SE-HPLC. In the case of fluorescent labeling, rhodamine-Mal (1% equivalent of total malyl groups) was conjugated with the polymer backbone. After conjugation was completed, excess of pyridyl dithiopropionate (PDP) was added to block remaining sulfhydryl groups (30 min, 20°C). The volume was adjusted to 4 mL by centrifugation over Vivaspin 20 and the product was purified over PD-10 column (in PBS). After removal of endotoxin using Triton X-114^17^, conjugates P/mPEG5000(2%)/LLL(40%)/a-CTLA-4(0.2%)/a-msTfR(0.2%) (P/a-CTLA-4) (Fig. 3a) and P/mPEG5000(2%)/LLL(40%)/a-PD-1(0.2%)/a-msTfR(0.2%) (P/a-PD-1) (Fig. 3b) were aliquoted and stored at -20°C.

### Size-exclusion HPLC

The synthesis of NICs were monitored with Hitachi HPLC system equipped with diode array detector (DAD) detectors (Hitachi, Pleasanton, CA) using size exclusion Polysep-GFC-P 4000 column (Phenomenex, Torrance, CA). The samples were monitored at 220, 260, and 280 nm with flow rate of 1 mL/min and PBS as eluent at 25°C.

### FTIR analysis

Solutions of antibody, pre-conjugate, and final nanoconjugates (P/a-CTLA-4 or P/a-PD-1) (1 mg/mL) in 100 μL PBS were lyophilized, and each mixed with 150 mg KBr. The mixtures were then analyzed using a Bruker alpha FTIR instrument fitted with a DRIFT module. An equivalent amount of PBS sample was measured and subtracted as background.

### Pull-down ELISA

NUNC MaxiSorp plates (Thermo Fisher Scientific, Waltham, MA) were coated with PD-1, CTLA-4 proteins (Acrobiosystems, Newark, DE), and mouse TfR (500 ng/well) (recombinant protein made by California Institute of Technology) in coating buffer (Protein Detector™ HRP Microwell Kit, SeraCare, Milford, MA) at 4°C overnight. The plates were blocked with 4% skim milk for 1 h at room temperature and washed once. The samples (a-CTLA-4, a-PD-1, a-msTfR, and nanoconjugates P/a-CTLA-4 or P/a-PD-1) were incubated in binding buffer containing 0.5% milk for 1 h followed by washing four times. Secondary HRP-labeled antibodies (goat anti-rat from Abcam, Cambridge, MA, goat anti-mouse and goat anti-hamster antibodies from SeraCare) were used for detection of free and conjugated a-msTfR and conjugated a-CTLA-4 or a-PD-1. The conjugated a-msTfR was detected with anti-rat/HRP secondary antibody when the other antibody a-CTLA-4 or a-PD-1 was attached to its plate-adsorbed antigen, to confirm the presence of both antibodies on one polymer chain (pull-down ELISA). Pull-down ELISA was also performed for detection of a-CTLA-4 or a-PD-1 when the other antibody a-msTfR was attached to its plate-adsorbed antigen similarly.

### Hydrodynamic diameter and ζ potential measurement

The NICs were characterized with respect to their size (hydrodynamic diameter) using a Malvern Zetasizer Nano (Malvern Instruments, UK). The diameter that is measured in DLS (Dynamic Light Scattering) refers to the particle diffusion within a fluid and is referred to as the hydrodynamic diameter corresponding to the diameter of a sphere that has the same translational diffusion coefficient as the NICs. All calculations were carried out using the Zetasizer 7.0 software. For the NIC size and ζ-potential measurements, the solutions were prepared in PBS at the concentration of 1-2 mg/mL after 0.2 μm membrane filtration. All copolymer solutions were prepared immediately before analysis at 25°C. Data represent the mean value obtained from three independent measurements.

### Malic acid assay

The content of PMLA in NICs was estimated by quantitative measurement of the malic acid produced after hydrolysis of NICs based on the photometric or fluorometric measurement of NADH formed by malate dehydrogenase (MDH)-catalyzed oxidation of L-malate to oxaloacetate. This enzyme is highly specific for L-malate, even in complex mixtures such as cellular extracts. Briefly, standard malic acid and sample solutions (100 μL) containing NICs were mixed with 100 μL of hydrochloric acid (12 M) and incubated for 16 h at 116°C in a 2 mL sealed glass vial. The cooled mixture was neutralized to near-neutral with 5 N NaOH equivalent to the mole of HCl. The standard malic acid and NIC hydrolysis solutions (20 μL) were mixed with 280 μL of glycine buffer (Sigma-Aldrich, St. Louis, MO), 20 μL of NAD+ (Sigma-Aldrich; 26.7 mg/mL, 40 mM), and 15 μL of L-malate-dehydrogenase (EMD Millipore, Temecula, CA; diluted 100-fold from commercial stock), allowed to stand for 30 min at 37°C, and the absorbance was read at 340 nm wavelength. The amount of malic acid in the sample was calculated from the standard curve.

### Cell line

Mouse glioblastoma cell line GL261 was a gift from Dr. Badies lab (UC San Diego, San Diego, CA) and was cultured in Dulbecco’s modified Eagle medium (DMEM) (ATCC, Manassas, VA) containing 10% fetal bovine serum with 1% penicillin (100 μg/mL), streptomycin (100 μg/mL), and amphotericin B (0.25 μg/mL) at 37°C with 5% CO_2_.

### Intracranial tumor model and treatment regimen

All animal experiments were performed with approval of Institutional Animal Care and Use Committee (IACUC) at Cedars-Sinai Medical Center (Los Angeles, CA).

Twenty thousand GL261 cells in 2 μL PBS were implanted intracranially into the right basal ganglia of immunocompetent 8-weeks old female C57BL/6J mice (The Jackson Laboratory, Sacramento, CA). All treatments were started on the sixth day after tumor cell inoculation. Free antibodies and NICs were administered at a dose of ~10 mg/kg via tail vein injections, twice per week for a total of 5 injections.

To prevent anaphylactic-like adverse effects, starting with the second treatment, all mice (including the control group) received 200 μg anti-histamine Triprolidine (Sigma-Aldrich) and 100 μg platelet-activating factor (PAF) antagonist CV6209 (Santa Cruz Biotechnology, Dallas, TX) via intraperitoneal injection, respectively, 30 min and 45 min prior to NIC injection.

### Immunostaining for drug delivery through BBB and T cell identification

For drug delivery experiments, tumor bearing mice alternatively injected with rhodamine-labeled NICs (P/a-CTLA-4 or P/a-PD-1) or rhodamine-labeled free mAbs (a-CTLA-4 or a-PD-1) were euthanized 4 h after injection. For T cell staining, tumor-bearing mice were alternatively treated with NICs (P/a-CTLA-4 or P/a-PD-1) or free antibodies (a-CTLA-4 or a-PD-1) and euthanized 24 h after the third treatment. In both drug delivery and T cell staining experiments, brains were embedded in OCT and sectioned using a Leica CM3050 S cryostat (Leica Biosystems, Buffalo Grove, IL). Tissue sections were air-dried at room temperature, fixed with 1% paraformaldehyde for 5 min and rinsed with PBS. Sections were blocked in 5% normal BSA and 0.1% Triton X-100 in PBS and incubated with anti-von Willebrand Factor (vWF) (ab11713, Abcam) labeled with AlexaFluor 488 or anti-Mouse CD8 Alpha (MCA609GA, Bio-Rad Laboratories, Hercules, CA), or anti-Mouse CD4 (MCA4635GA, Bio-Rad) and anti-Mouse FoxP3 (ab20034, Abcam). As secondary antibodies we used goat anti-rat (FITC) (112-095-167), goat anti-mouse (TRITC) (115-025-166), and donkey anti-rat (TRITC) (712025153), all from Jackson Immunoresearch. Sections were mounted with ProLongGold Antifade (Thermo Fisher Scientific) mounting medium containing 4’,6-diamidino-2-phenylindole (DAPI) to counterstain cell nuclei. Images were captured using a Leica DM6000 B microscope (Leica).

Optical imaging data were analyzed using FIJI software^51^. Images acquired at 40X or 63X magnification were first normalized for size. This was done by cropping the 40X images (85% of data) to a dimension that equaled the 63X image. The position of the crop was randomized so that cells included in the final analysis were objectively selected. Total cell numbers were calculated from manual counts of DAPI-labeled nuclei; counts were performed with the FIJI cell counter tool. Cells that were labeled with anti-CD8, anti-CD4 and anti-FoxP3 antibodies were counted separately by two investigators, and ratios were calculated against the number of DAPI-labeled cells. We analyzed 3-5 images from 2 mice for each drug condition. In total, 38 images were analyzed to produce CD8+ data, and 43 images were analyzed to produce FoxP3+ cell counts. Statistical analysis and plots were made using Prism 6.0 (GraphPad, La Jolla, CA). Cell counts were compared by one-way ANOVA combined with pairwise post-hoc comparisons of individual data points; exact parameters and tests are separately indicated for each result.

### Flow cytometry

Tumors were harvested at euthanasia. Tissue samples were chopped and incubated in RPMI medium (ATCC) containing 1.67 U/mL Liberase TL (Roche Diagnostics GmbH, Mannheim, Germany) for 30 min at 37°C and mashed through a 70 μm strainer to obtain single cell suspension. Cell samples were blocked with rat anti-mouse CD16/32 (Mouse BD Fc Block, clone 2.4G2, BD Biosciences) and then stained with BV421 rat anti-mouse CD3 molecular complex (BD Biosciences), APC/Cy7 anti-mouse CD4 (Biolegend, San Diego, CA) and PE-Texas Red rat anti-mouse CD8α (Thermo Fisher Scientific). Cells were permeabilized with FoxP3 Staining Buffer Set (Affimetrix, San Diego, CA) and stained with anti-mouse/rat FoxP3 FITC (clone FJK-16s) (Affimetrix). Cells were then acquired using a BD LSR Fortessa cell analyzer equipped with BD FACSDiva software (BD Biosciences). Data were analyzed using FlowJo software (FlowJo, Ashland, OR).

### Cytokine analysis

Blood was drawn from tumor bearing mice injected with NICs 24 h after the third treatment, and serum was separated for cytokine analysis. The multiplex cytokine assay was performed using a custom Bio-Plex Pro Assay kit (Bio-Rad) including antibodies targeting IL-1β, IL-2, IL-4, IL-5, IL-6, IL-10, IL-12(p70), IFN-γ, and TNF-α, according to manufacturer’s instructions. Results were obtained using a Bio-Plex 200 System (Bio-Rad) equipped with Bio-Plex Manager Software (Bio-Rad) and data were processed using the same software.

### Statistical analysis

Statistical analysis of survival data was carried out using Kaplan-Meier curves and log-rank test by GraphPad Prism 6.0 software (GraphPad). Multiplex cytokine assay and flow cytometry results were presented as mean ± SEM and multiple t-test analyses or ANOVA were used to compare treatment groups. P<0.05 was considered statistically significant.

## Acknowledgments

This work was supported by NIH R01 Grants EY013431 (AVL), CA 206220-01 and CA 230858-01 (JL).

## Author contributions

A.G. and A.C. performed experiments and data analysis, and wrote the manuscript; K.L.B., E.H., A.V.L., A.J.K., and M.L.P. contributed ideas and commented on the manuscript; T.S., E.S., and V.A.L., J. M. assisted with in vivo work; L.L.I. assisted with drug characterization; O.B. performed optical imaging data analysis; R.P. performed antibody labelling for immunostaining; Z.G. produced PMLA; H.D. performed drug synthesis and characterization; H.D., E.H, A.V.L. and J.Y.L., conceived and designed the experiments and wrote the manuscript.

## Survival of intracranial glioma bearing mice after treatment with nano immunoconjugates

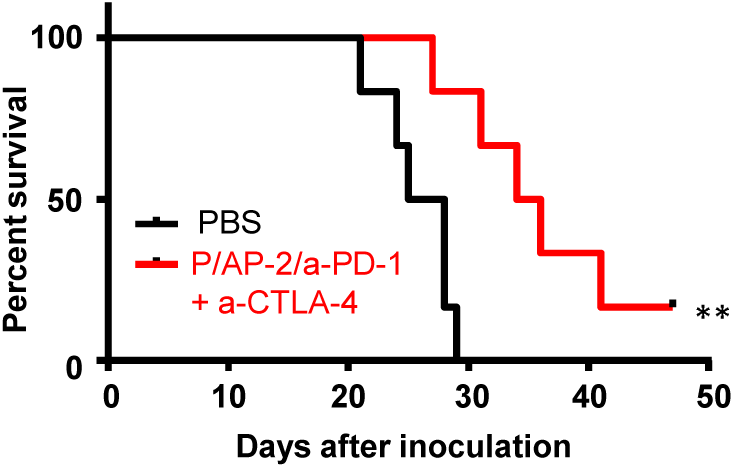
Kaplan-Meier survival plot after the treatment with nano immunoconjugates. Combination of P/CTLA-4+P/PD-1 significantly increases survival vs. PBS treated mice: with peptide p<0.006.

## References

1. Fontana F, Liu D, Hirvonen J, Santos HA. Delivery of therapeutics with nanoparticles: what’s new in cancer immunotherapy? Wiley Interdiscip Rev Nanomed Nanobiotechnol 2017, 9(1).

2. Hodi FS, O’Day SJ, McDermott DF, Weber RW, Sosman JA, Haanen JB, et al. Improved survival with ipilimumab in patients with metastatic melanoma. N Engl J Med 2010, 363(8): 711–723.

3. Robert C, Thomas L, Bondarenko I, O’Day S, Weber J, Garbe C, et al. Ipilimumab plus dacarbazine for previously untreated metastatic melanoma. N Engl J Med 2011, 364(26): 2517–2526.

4. Wainwright DA, Chang AL, Dey M, Balyasnikova IV, Kim CK, Tobias A, et al. Durable therapeutic efficacy utilizing combinatorial blockade against IDO, CTLA-4, and PD-L1 in mice with brain tumors. Clin Cancer Res 2014, 20(20): 5290–5301.

5. Belcaid Z, Phallen JA, Zeng J, See AP, Mathios D, Gottschalk C, et al. Focal radiation therapy combined with 4-1BB activation and CTLA-4 blockade yields long-term survival and a protective antigen-specific memory response in a murine glioma model. PLoS One 2014, 9(7): e101764.

6. Fecci PE, Heimberger AB, Sampson JH. Immunotherapy for primary brain tumors: no longer a matter of privilege. Clin Cancer Res 2014, 20(22): 5620–5629.

7. Sampson JH, Vlahovic G, Sahebjam S, Omuro AMP, Baehring JM, Hafler DA, et al. Preliminary safety and activity of nivolumab and its combination with ipilimumab in recurrent glioblastoma (GBM): CHECKMATE-143. J Clin Oncol 2015, 33(15).

8. Pardridge WM. Drug transport across the blood-brain barrier. J Cereb Blood Flow Metab 2012, 32(11): 1959–1972.

9. Villasenor R, Ozmen L, Messaddeq N, Gruninger F, Loetscher H, Keller A, et al. Trafficking of Endogenous Immunoglobulins by Endothelial Cells at the Blood-Brain Barrier. Sci Rep 2016, 6: 25658.

10. St-Amour I, Pare I, Alata W, Coulombe K, Ringuette-Goulet C, Drouin-Ouellet J, et al. Brain bioavailability of human intravenous immunoglobulin and its transport through the murine blood-brain barrier. J Cereb Blood Flow Metab 2013, 33(12): 1983–1992.

11. Lim M, Xia Y, Bettegowda C, Weller M. Current state of immunotherapy for glioblastoma. Nat Rev Clin Oncol 2018, 15(7): 422–442.

12. Ljubimova JY, Ding H, Portilla-Arias J, Patil R, Gangalum PR, Chesnokova A, et al. Polymalic acid-based nano biopolymers for targeting of multiple tumor markers: an opportunity for personalized medicine? J Vis Exp 2014(88).

13. Patil R, Ljubimov AV, Gangalum PR, Ding H, Portilla-Arias J, Wagner S, et al. MRI virtual biopsy and treatment of brain metastatic tumors with targeted nanobioconjugates: nanoclinic in the brain. ACS Nano 2015, 9(5): 5594–5608.

14. Lee B-S, Vert M, Holler E. Water-soluble Aliphatic Polyesters: Poly(malic acid)s. Biopolymers Online. Wiley-VCH Verlag GmbH & Co. KGaA, 2005.

15. Ding H, Inoue S, Ljubimov AV, Patil R, Portilla-Arias J, Hu J, et al. Inhibition of brain tumor growth by intravenous poly (beta-L-malic acid) nanobioconjugate with pH-dependent drug release [corrected]. Proc Natl Acad Sci U S A 2010, 107(42): 18143–18148.

16. Ding H, Portilla-Arias J, Patil R, Black KL, Ljubimova JY, Holler E. Distinct mechanisms of membrane permeation induced by two polymalic acid copolymers. Biomaterials 2013, 34(1): 217–225.

17. Aida Y, Pabst MJ. Removal of Endotoxin from Protein Solutions by Phase-Separation Using Triton X-114. J Immunol Methods 1990, 132(2): 191–195.

18. Jure-Kunkel MN, Masters G, Girit E, Dito G, Lee FY. Antitumor activity of anti-CTLA-4 monoclonal antibody (mAb) in combination with ixabepilone in preclinical tumor models. J Clin Oncol 2008, 26(15).

19. Wing K, Onishi Y, Prieto-Martin P, Yamaguchi T, Miyara M, Fehervari Z, et al. CTLA-4 control over Foxp3+ regulatory T cell function. Science 2008, 322(5899): 271–275.

20. Jonsson F, Mancardi DA, Kita Y, Karasuyama H, Iannascoli B, Van Rooijen N, et al. Mouse and human neutrophils induce anaphylaxis. J Clin Invest 2011, 121(4): 1484–1496.

21. Khodoun MV, Strait R, Armstrong L, Yanase N, Finkelman FD. Identification of markers that distinguish IgE-from IgG-mediated anaphylaxis. P Natl Acad Sci USA 2011, 108(30): 12413–12418.

22. Murphy JT, Burey AP, Beebe AM, Gu D, Presta LG, Merghoub T, et al. Anaphylaxis caused by repetitive doses of a GITR agonist monoclonal antibody in mice. Blood 2014, 123(14): 2172–2180.

23. [cited]Available from: https://www.cancer.gov/research/key-initiatives/moonshot-cancer-initiative/funding?cid=eb_govdel_en_grantees_moonshot-foa_announcement_rsr.

24. Ribas A, Shin DS, Zaretsky J, Frederiksen J, Cornish A, Avramis E, et al. PD-1 Blockade Expands Intratumoral Memory T Cells. Cancer Immunol Res 2016, 4(3): 194–203.

25. Ribas A. Anti-CTLA4 Antibody Clinical Trials in Melanoma. Update Cancer Ther 2007, 2(3): 133–139.

26. Reardon DA, Gokhale PC, Klein SR, Ligon KL, Rodig SJ, Ramkissoon SH, et al. Glioblastoma Eradication Following Immune Checkpoint Blockade in an Orthotopic, Immunocompetent Model. Cancer Immunol Res 2016, 4(2): 124–135.

27. Martinez-Lostao L, Anel A, Pardo J. How Do Cytotoxic Lymphocytes Kill Cancer Cells? Clin Cancer Res 2015, 21(22): 5047–5056.

28. Bayer AL, Pugliese A, Malek TR. The IL-2/IL-2R system: from basic science to therapeutic applications to enhance immune regulation. Immunol Res 2013, 57(1-3): 197–209.

29. Geng X, Zhang R, Yang G, Jiang W, Xu C. Interleukin-2 and autoimmune disease occurrence and therapy. Eur Rev Med Pharmacol Sci 2012, 16(11): 1462–1467.

30. Morgan DA, Ruscetti FW, Gallo R. Selective in vitro growth of T lymphocytes from normal human bone marrows. Science 1976, 193(4257): 1007–1008.

31. Aloisi F. Immune function of microglia. Glia 2001, 36(2): 165–179.

32. Hanisch UK. Microglia as a source and target of cytokines. Glia 2002, 40(2): 140–155.

33. Gerger A, Renner W, Langsenlehner T, Hofmann G, Knechtel G, Szkandera J, et al. Association of interleukin-10 gene variation with breast cancer prognosis. Breast Cancer Res Tr 2010, 119(3): 701–705.

34. Tabrez S, Ali M, Jabir NR, Firoz CK, Ashraf GM, Hindawi S, et al. A Putative Association of Interleukin-10 Promoter Polymorphisms with Cardiovascular Disease. Iubmb Life 2017, 69(7): 522–527.

35. Haabeth OAW, Lorvik KB, Yagita H, Bogen B, Corthay A. Interleukin-1 is required for cancer eradication mediated by tumor-specific Th1 cells. Oncoimmunology 2016, 5(1).

36. Kim ES, Choi YE, Hwang SJ, Han YH, Park MJ, Bae IH. IL-4, a direct target of miR-340/429, is involved in radiation-induced aggressive tumor behavior in human carcinoma cells. Oncotarget 2016, 7(52): 86836–86856.

37. Rand RW, Kreitman RJ, Patronas N, Varricchio F, Pastan I, Puri RK. Intratumoral administration of recombinant circularly permuted interleukin-4-Pseudomonas exotoxin in patients with high-grade glioma. Clinical Cancer Research 2000, 6(6): 2157–2165.

38. Markert JM, Cody JJ, Parker JN, Coleman JM, Price KH, Kern ER, et al. Preclinical Evaluation of a Genetically Engineered Herpes Simplex Virus Expressing Interleukin-12. J Virol 2012, 86(9): 5304–5313.

39. Chu MB, Fesler MJ, Armbrecht ES, Fosko SW, Hsueh E, Richart JM. High-Dose Interleukin-2 (HD IL-2) Therapy Should Be Considered for Treatment of Patients with Melanoma Brain Metastases. Chemother Res Pract 2013, 2013: 726925.

40. Powell S, Dudek AZ. Single-institution outcome of high-dose interleukin-2 (HD IL-2) therapy for metastatic melanoma and analysis of favorable response in brain metastases. Anticancer Res 2009, 29(10): 4189–4193.

41. Atkins MB, Clark JI, Quinn DI. Immune checkpoint inhibitors in advanced renal cell carcinoma: experience to date and future directions. Ann Oncol 2017, 28(7): 1484–1494.

42. Gadani SP, Cronk JC, Norris GT, Kipnis J. IL-4 in the brain: a cytokine to remember. J Immunol 2012, 189(9): 4213–4219.

43. Lamichhane P, Karyampudi L, Shreeder B, Krempski J, Bahr D, Daum J, et al. IL10 Release upon PD-1 Blockade Sustains Immunosuppression in Ovarian Cancer. Cancer Res 2017, 77(23): 6667–6678.

44. Redmond WL, Linch SN, Kasiewicz MJ. Combined Targeting of Costimulatory (OX40) and Coinhibitory (CTLA-4) Pathways Elicits Potent Effector T Cells Capable of Driving Robust Antitumor Immunity. Cancer Immunology Research 2014, 2(2): 142–153.

45. Galli SJ. Pathogenesis and management of anaphylaxis: current status and future challenges. J Allergy Clin Immunol 2005, 115(3): 571–574.

46. Kraft S, Kinet JP. New developments in FcepsilonRI regulation, function and inhibition. Nat Rev Immunol 2007, 7(5): 365–378.

47. Sharpe AH, Pauken KE. The diverse functions of the PD1 inhibitory pathway. Nat Rev Immunol 2018, 18(3): 153–167.

48. Preusser M, Winkler F, Valiente M, Manegold C, Moyal E, Widhalm G, et al. Recent advances in the biology and treatment of brain metastases of non-small cell lung cancer: summary of a multidisciplinary roundtable discussion. ESMO Open 2018, 3(1): e000262.

49. Owonikoko TK, Arbiser J, Zelnak A, Shu HKG, Shim H, Robin AM, et al. Current approaches to the treatment of metastatic brain tumours. Nature Reviews Clinical Oncology 2014, 11(4): 203–222.

50. Lee BS, Holler E. beta-poly(L-malate) production by non-growing microplasmodia of Physarum polycephalum -Effects of metabolic intermediates and inhibitors. Fems Microbiol Lett 2000, 193(1): 69–74.

51. Schindelin J, Arganda-Carreras I, Frise E, Kaynig V, Longair M, Pietzsch T, et al. Fiji: an open-source platform for biological-image analysis. Nat Methods 2012, 9(7): 676–682.

